# Treatment with the ribosome biogenesis inhibitor CX-5461 increases platelet count in humans and enhances murine megakaryopoiesis

**DOI:** 10.1101/2025.06.15.659730

**Authors:** Vijay Bhoopalan, Amandeep Kaur, James I. Hearn, Kylee H. Maclachlan, Lixinyu Liu, Rita Ferreira, Sidra A. Ali, Yee Lin Thong, Nadine Hein, Samina Nazir, Simone A. Brysland, Si Ming Man, Simon J. Harrison, Robert K. Andrews, Koji Eto, Philip Y-I Choi, Jiayu Wen, Kate M. Hannan, Ross D. Hannan, Elizabeth E. Gardiner

**Affiliations:** Genome Sciences and Cancer Division, The John Curtin School of Medical Research, The Australian National University, Canberra, Australia; Memorial Sloan Kettering Cancer Centre, New York, United States of America; Immunology and Infectious Diseases Division, The John Curtin School of Medical Research, The Australian National University, Canberra, Australia; Sir Peter MacCallum Department of Oncology, University of Melbourne, Melbourne, Australia; Center for iPS Cell Research and Application, Kyoto University, Japan; The Canberra Hospital, Australian Capital Territory, Australia

## Abstract

Thrombocytopenia is a common and serious complication of anticancer therapies. Here, we identify a novel thrombopoietic activity of the first-in-class ribosome biogenesis inhibitor CX-5461. In a phase I trial, 56% (9/16) of patients exhibited up to a 34% increase in platelet count following a single dose of CX-5461. In mice, CX-5461 elicited a rapid, reversible, and sustained ∼1.7-fold increase in platelet numbers without altering platelet function, lifespan, or inflammatory cytokines. Bone marrow analysis revealed a specific expansion of megakaryocytes (MKs), increased Sca1⁺ MKs, and selective enrichment of MK-biased multipotent progenitor 2, independent of thrombopoietin (TPO) or c-mpl signalling. CX-5461 also mitigated carboplatin-induced thrombocytopenia, accelerating platelet recovery. Single-cell RNA sequencing and RNA velocity analysis confirmed enhanced differentiation of MK progenitors. These findings demonstrate that inhibition of ribosome biogenesis promotes TPO-independent megakaryopoiesis and identifies a previously unrecognised therapeutic opportunity to support platelet recovery in cancer treatment and potentially other thrombocytopenic states.

## Introduction

Platelets play a crucial role in maintaining haemostasis, facilitating clot formation to prevent bleeding following vessel injury. Reduced levels of circulating platelets (thrombocytopenia) elevate bleeding potential and interfere with innate immune responses.^1^ Thrombocytopenia is a comorbidity frequently occurring in disease and as a side effect of clinical therapies, such as cancer chemotherapy.^2^ Treatments that directly increase platelet count are limited to thrombopoietin (TPO) receptor agonists and platelet transfusion, which carry potential side effects and are not without risk.^3^ TPO mimetics bind to the myeloproliferative leukemia protein (c-mpl) receptor, which is expressed on the surface of haematopoietic stem cells (HSCs) and megakaryocytes (MKs).^4^ Activation of c-mpl receptor signalling accelerates HSC self-renewal and drives differentiation of HSCs along myeloid pathways, forming megakaryocytic-erythroid progenitors (MEPs) and MKs, enhancing MK maturation and platelet production.^1,2^ However, platelet biogenesis initiated by TPO mimetics is slow and associated with myelofibrosis.^5^ There is a critical unmet clinical need for agents that stimulate thrombopoiesis through TPO-independent mechanisms to safely and effectively restore platelet counts in thrombocytopenic patients.^4^

CX-5461 is a first-generation inhibitor of ribosome biogenesis that inhibits RNA polymerase I (Pol I)-mediated transcription of ribosomal genes (rDNA) to produce ribosomal RNAs (rRNA) that form the nucleic backbone of ribosomes that translate messenger RNA (mRNA) into proteins. CX-5461 inhibits Pol I transcription by preventing the recruitment of selectivity-factor 1, a Pol I-specific transcription factor important for preinitiation complex formation.^6^ When rRNA synthesis is downregulated, this disrupts proliferative capacity and promotes lineage commitment and cellular differentiation.^7,8^

CX-5461 (pidnarulex™) has completed phase I clinical trials in haematological malignancies and solid tumour patients, and was shown to be well-tolerated. Palmar-plantar erythrodysesthesia photosensitivity was a dose-independent adverse event.^9,10^ In the current study, we report that patients treated with CX-5461 show increased platelet counts. Moreover, mice treated with CX-5461 for up to 42 days show a rapid and sustained increase in platelet counts with normal platelet function and lifespan. Our study provides the first definitive evidence of drug-induced robust, TPO-independent thrombopoiesis, resulting in a sustained increase in functionally competent circulating platelets in both mice and humans. This unique property distinguishes CX-5461, and likely other inhibitors of RNA Pol I transcription^11^ from other anticancer agents and highlights its potential dual utility as both an antineoplastic and a thrombopoietic agent, particularly in settings of chemotherapy-induced thrombocytopenia or in patients with impaired TPO signalling.

## Results

### Changes to platelet count in patients with haematological malignancy in receipt of CX-5461

Details of study design, patient characteristics and outcomes from a phase I dose escalation study of CX-5461 in patients with haematological cancer have been reported.^9^ Sixteen patients were treated with CX-5461 in sequential cohorts at 25-250 mg/m^2^ (**Table 1).** Anaemia and mild decreases in neutrophil and platelet counts were noted in some patients.^9^ On closer scrutiny of the haematological data, 9/16 (56%) of patients treated with a single dose of CX-5461 showed increases (1-34%) in platelet count at day 15 post treatment. Platelet count increments were most frequently observed in people who received a higher dose of CX-5461 **(Table 1, STable 1)** and commenced from day 8 in responders. To assess whether CX-5461 exposure had triggered platelet activation, we evaluated plasma for levels of soluble GPVI (sGPVI), a platelet-specific activation marker. GPVI is stable on resting platelets, but sGPVI can be metalloproteolytically released from activated platelets in various pathologies.^12–14^

**Table 1:** Variation in platelet counts in patients receiving a single dose of CX-. 5461. Patients with haematological cancer received the indicated dose of CX-5461 as part of a phase I dose-escalation study.^9^ Platelet counts remained within reference ranges (100-450 x 10^9^/L) for all patients except PMC-12. 9/16 (56%) patients had increased platelet counts on day 15 (red values).

| Trial ID | Gender | Cancer | CX-5461 dose (mg/m<sup>2</sup>) | % Change in platelet counts over baseline |  |  |  |
| --- | --- | --- | --- | --- | --- | --- | --- |
|  |  |  |  | Day 2 | Day 3 | Day 8 | Day 15 |
| PMC-01 | M | MM | 25 | 1.30 | 0.65 | -1.95 | 1.30 |
| PMC-03 | F | MM | 25 | -12.98 | -3.85 | 26.44 | 15.38 |
| PMC-04 | F | FL->DLBCL | 25 | -19.03 | NA | -34.60 | -21.11 |
| PMC-05 | F | CTCL | 50 | -26.69 | -20.95 | -4.73 | 12.16 |
| PMC-06 | F | HL | 50 | -10.10 | -8.65 | -43.27 | -64.42 |
| PMC-07 | M | CLL->DLBCL | 50 | -5.05 | -19.19 | -40.40 | -66.67 |
| PMC-08 | F | DLBCL | 50 | -5.08 | -1.69 | 12.99 | 14.69 |
| PMC-09 | M | MM | 100 | -14.38 | -18.75 | -25.63 | -59.38 |
| PMC-10 | M | T-PLL | 100 | -14.08 | -4.37 | -26.70 | -92.72 |
| PMC-11 | F | DLBCL | 100 | -6.38 | -10.11 | NA | -20.21 |
| PMC-12 | M | HL | 100 | -1.33 | 11.41 | -1.52 | 15.02 |
| PMC-13 | F | MM | 170 | 0.97 | 5.83 | 6.31 | 11.65 |
| PMC-14 | M | MM | 170 | -9.56 | -4.41 | -13.24 | -4.41 |
| PMC-15 | F | DLBCL | 170 | 28.47 | 26.28 | 40.88 | 34.31 |
| PMC-16 | F | MM | 250 | -5.08 | -5.93 | -1.69 | 5.93 |
| PMC-17 | M | ALCL | 250 | 12.43 | 7.10 | 16.57 | 23.08 |

*Multiple myeloma (MM), Follicular lymphoma (FL), diffuse large B cell lymphoma (DLBCL),* *cutaneous T-cell lymphoma (CTCL), chronic lymphocytic leukemia (CLL), Hodgkin's* *lymphoma (HL), T-cell prolymphocytic leukemia (T-PLL), Anaplastic large cell lymphoma* *(ALCL). Not available (NA).*

Levels of sGPVI in plasma from CX-5461-treated patients were significantly reduced compared to healthy controls; however, no significant differences were observed within the patient group pre- and post-CX-5461 treatment **Extended data (ED) 1A),** suggesting that CX-5461 treatment did not induce any peripheral platelet activation. Levels of plasma cytokines largely remained within normal ranges **(ED. 1B)**.

### CX-5461-treated mice showed elevated platelet counts and immature (reticulated) platelet fractions

While CX-5461 therapy achieved strong therapeutic responses in murine leukaemia models,^15^ the impact on platelet counts was not examined in these studies. To explore whether CX-5461 treatment in mice recapitulated the observed increase in platelet counts seen in the clinical trial, we treated mice with 10-35 mg/kg of CX-5461 or vehicle 3 times a week for up to 42 days. CX-5461 treatment increased circulating platelet count commencing at day 7 and was maintained for the duration of CX-5461 treatment (**Fig. 1A, B**). The dose-dependent increase was specific to platelets, enumerated using either an automated analyser (**Fig. 1B**) or by flow cytometry (**Fig. 1A and ED. 2A**). Red blood cells (RBC) and lymphocytes (**EDs. 2B and C**) were either reduced or remained unchanged throughout the treatment period. Upon treatment discontinuation, platelets returned to normal pretreatment levels within 7 days (**Fig. 1C**). Reticulated (immature) platelets were elevated 2-fold in mice treated with 35 mg/kg CX-5461 (**Fig. 1D**). This data suggests that CX-5461 treatment induces a rapid, sustained and isolated increase in the platelet count by promoting platelet biogenesis. A dosing regimen of 35 mg/kg CX-5461 administered three times per week was selected for subsequent studies based on prior pharmacokinetic and tolerability data supporting its translational relevance. Similar results were obtained using a second RNA polymerase I inhibitor, BMH-21 (thrice weekly injections of 75 mg/kg; **ED. 2D**), indicating that the thrombopoietic effect was not unique to CX-5461 but could be recapitulated using another RNA Pol I inhibitor. Moreover, BMH-21 has not been reported to inhibit Topo II, ruling out the latter as the mechanism mediating platelet levels

**Figure 1:**
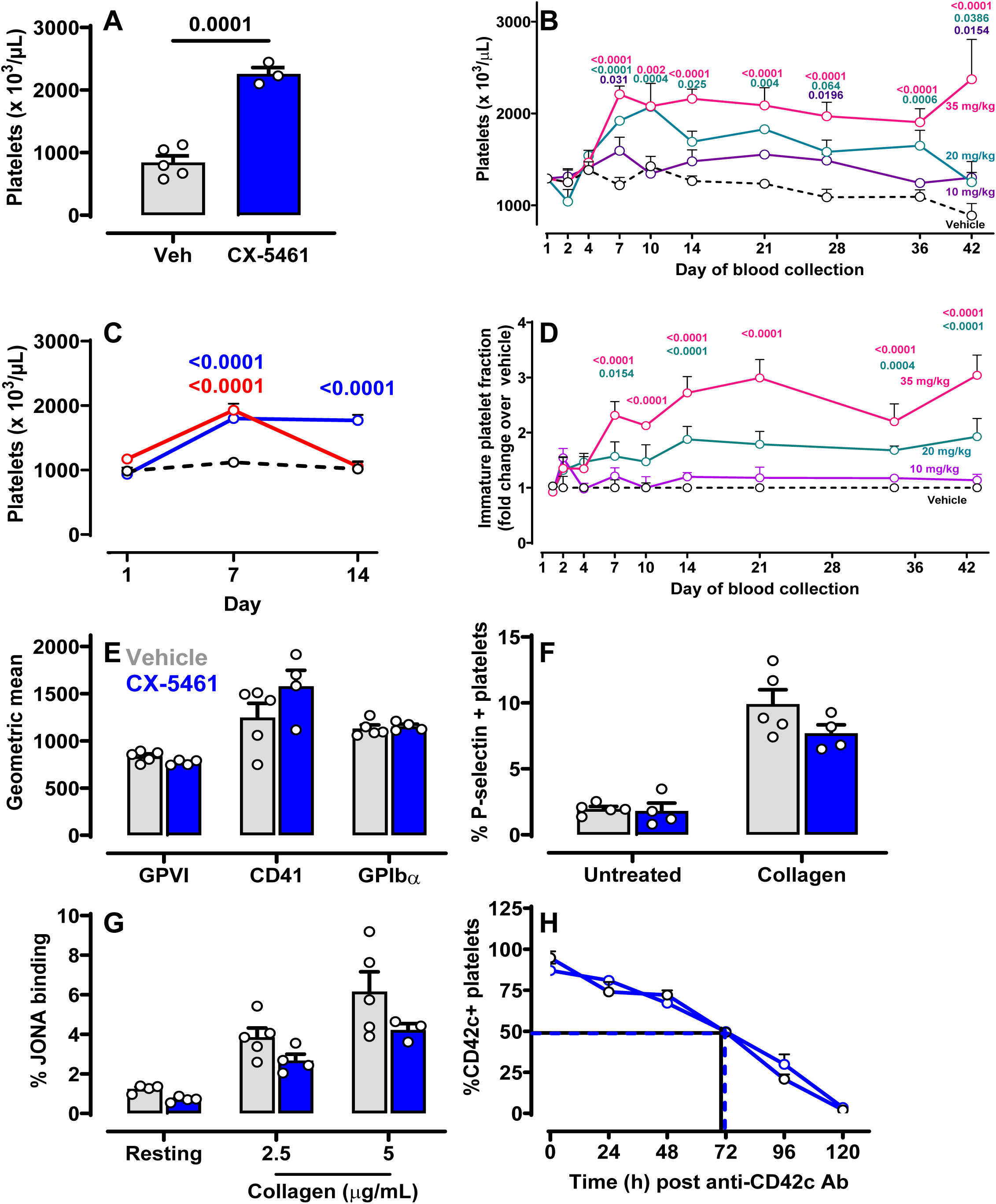
CX-5461 treated mice show increased peripheral platelet count without altering platelet function. C57BL/6 mice were randomly assigned to four groups and treated with either vehicle (control) or CX-5461 at 10, 20 or 35 mg/kg (3 doses/week), respectively, for 42 days. Tail vein blood samples were collected to quantify **(A)** platelets on day 14 post-CX-5461 (35 mg/kg) treatment using counting beads in a flow cytometer. **(B)** Peripheral platelet counts in mice treated with vehicle or 10, 20 or 35 mg/kg CX-5461. **(C)** Peripheral platelet counts in mice treated with vehicle (Black) or 35 mg/kg CX-5461 for 7 (red) or 14 (blue) days. All subsequent experimental data was collected from mice treated with 35 mg/kg CX-5461 or vehicle at day 42. **(D)** Thiazole orange-stained reticulated (immature) platelet fraction. **(E)** Receptor levels on circulating platelets. **(F)** P-selectin levels on untreated or 10 µg/mL collagen-activated platelets. **(G)** Activated αIIbβ3 levels (JON/A antibody binding) on resting and 2.5 or 5 μg/ml collagen-treated platelets. **(H)** Survival of Dylight 488-conjugated CD42c-labelled platelets in mice treated with vehicle (black) or CX-5461 (blue). The baseline (100% GPIbβ^+^ platelets) was set at 1 h post-injection, and % of labelled platelets was monitored over time by flow cytometry (n=3-6 mice/group). Panels A-G n= 3-8 mice/group were analysed. The significance among groups was analysed by student’s t-test (panel A), two-way ANOVA with Sidak’s multiple comparisons tests. Statistically significant numerical p-values are presented on the graphs.

### Platelets from CX-5461-treated mice exhibited normal receptor levels, function and circulating half-life

Accelerated platelet production can lead to altered platelet receptor levels and responsiveness to ligands.^16^ Treatment with CX-5461 for 42 days did not significantly alter the levels of platelet-specific surface receptors GPVI, αIIb integrin subunit or GPIbα compared to vehicle-treated mice (**Fig. 1E**). Enhanced platelet activation was not evident as levels of granule protein P-selectin and active αIIbβ3 (JON/A binding) on resting platelets were similar between treatment groups. Furthermore, platelets from CX-5461- or vehicle-treated mice displayed normal responses to collagen exposure (**Fig. 1F, G**). There was no significant change in the half-life of circulating platelets between the two groups, suggesting that alterations to platelet clearance did not contribute to the observed increase in circulating platelets (**Fig. 1H**). Reductions in circulating RBC (**ED. 2B**) and lymphocyte (**ED. 2C**) numbers were observed along with mild reductions in some other cell types (**STable 2**) in CX-5461-treated mice, however the RBC half-life remained unaffected (**ED. 2E**), indicating that the decrease in RBCs was due to reduced production rather than accelerated clearance. Together, these data suggest that CX-5461 treatment increased numbers of functionally normal platelets, which circulated with a normal half-life, whilst RBC and lymphocyte levels were diminished.

### CX-5461 treatment alters the cell composition within the BM

As CX-5461 treatment accelerated platelet biogenesis, we performed a detailed analysis of BM cellularity. The number of MKs was significantly higher in BM from CX-5461-treated mice compared to vehicle-treated mice on day 14, by histological **(Fig. 2Ai, Aii)** or flow cytometric analysis of immunostained FSC^high^ CD41^+^ cells at day 14 and 42 **(Fig. 2B, ED. 3A)**. Elevated MK numbers coincided with the increase in platelet count **(Fig. 2B)**. Total BM cellularity was unaffected by CX-5461 treatment, suggesting the increased frequency of MK was due to enhanced megakaryopoiesis (**Fig. 2C)**. MK ploidy showed no significant differences between treatment groups, while CX-5461-treatment increased CD42b expression is observed in high ploidy (16N, 32N and 128N) MKs (**Fig. 2D).** Similarly, proplatelet formation assays *ex vivo*, using BM isolated from control or CX-5461-treated mice showed no differences in MK potential to form proplatelets **(Fig. 2E)**.

**Figure 2:**
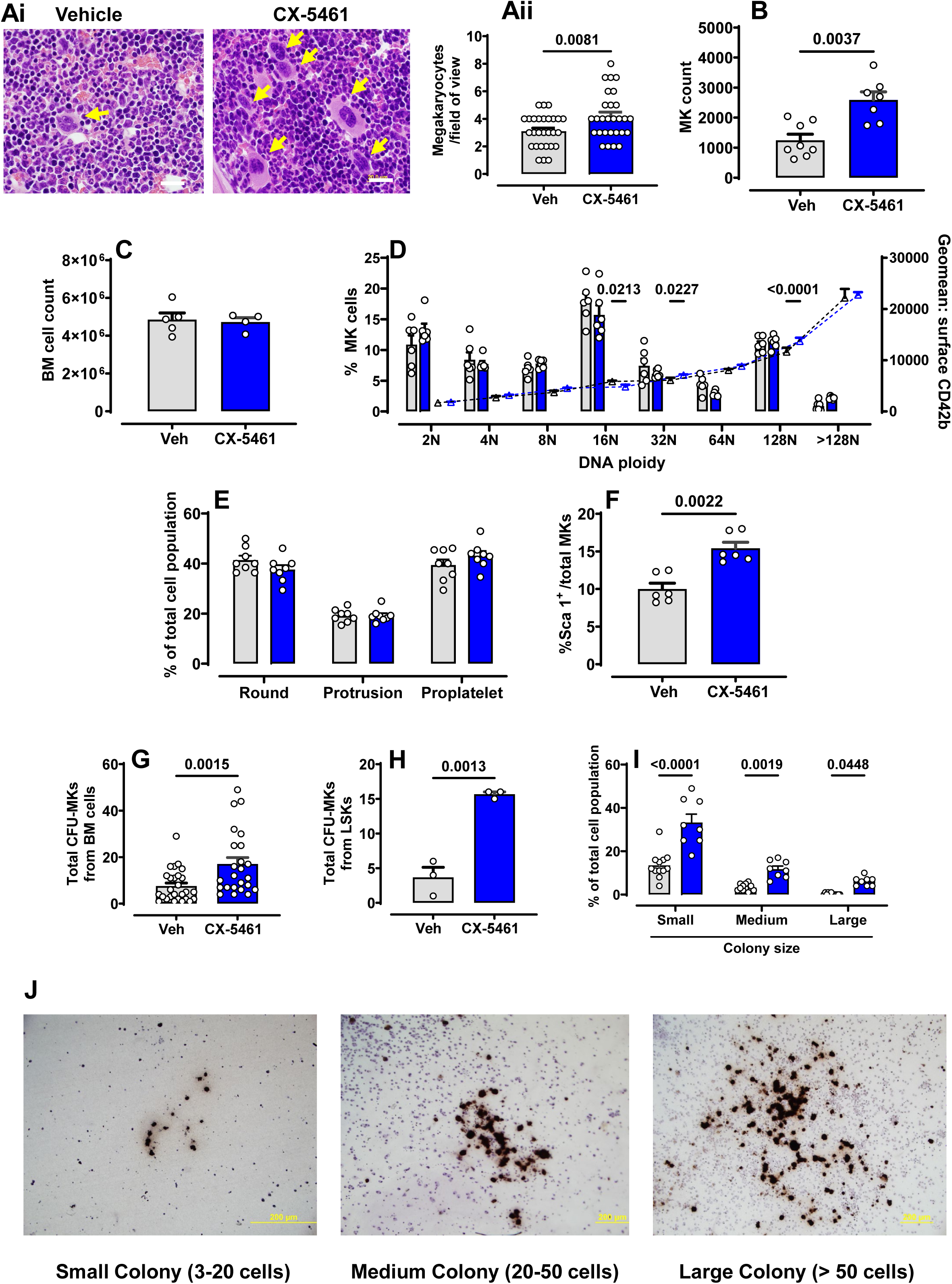
Megakaryocyte population increases in mice treated with CX-5461. C57BL/6 mice were treated with 35 mg/kg CX-5461 or vehicle for 14 days. **(A) (i)** H&E staining of BM sections at 100X magnification; yellow arrows indicate MKs, scale bar=20 µm **(ii)** MKs were manually counted from n=9-10 fields of view per treatment/mice from n=3 mice/group (vehicle or CX-5461). BM was isolated, and MK analyses were performed by flow cytometry **(B)** Numbers of CD41^+^CD42b^+^ MKs were measured in 10^6^ BM cell events. **(C)** Total BM cellularity **(D)** MK ploidy quantified using PI (bar graphs) and maturation quantified using CD42b^+^ (triangular symbols and line graphs) from vehicle-(black) or CX-5461-treated (blue) mice by flow cytometry. **(E)** Analysis of round, protrusion forming and proplatelet forming MKs in BM cultures *ex vivo*. **(F)** Percentages of Sca1^+^ and CD41^+^ double-positive cells in the total BM MK population were measured (n=6). **(G)** Graph showing the number of MK colonies on MegaCult assay plate arising from total BM (n=8 mice) or **(H)** sorted LSK cell populations (n=3 mice). The colonies were counted and scored as described previously.^49^ **(I)** Bar graph showing stratification of MK colonies evaluated based on their size, categorized as either small (3-20), medium (20-30) or large (>50 cells) (n=8 mice) **(J)** Representative images of BM cells cultured on MegaCult at day 7. MKs were stained using acetylcholinesterase (brown colour cells). Data are represented as mean ± SEM. The significance among groups was analysed using either a student’s t-test (panels Aii, F, and H), Mann Whitney’s test (panels B, C, and G) or two-way ANOVA with Sidak’s (panels D-E and I) multiple comparisons test. Statistically significant numerical p-values are presented on the graphs.

To further investigate CX-5461-induced changes, we characterised MK surface markers, focusing on Sca1, as elevated Sca1⁺ MKs have been reported under haematopoietic stress.^17^ We assessed Sca1 levels within the total MK cell population (CD41⁺/FSC^high^) and found that, while vehicle-treated mice exhibited Sca1⁺ MK proportions consistent with previously reported levels⁹, CX-5461-treated BM showed a significant increase in Sca1⁺ MK frequency **(Fig. 2F, ED. 3B)**.

To evaluate whether BM from CX-5461-treated mice directed differentiation towards the MK lineage, we performed MK colony-forming assays. Whole BM and flow cytometrically sorted lineage-negative (Lin^-^) Sca1^+^cKit^+^ (LSK) cells from control and CX-5461-treated mice were cultured for 8 days in a medium containing TPO, IL-3, IL-6 and IL-11. MK colonies identified by acetylcholine esterase activity were categorized as small (0-20 cells), medium (20-50 cells) and large (>50 cells) colonies and enumerated. Both total BM and LSK cells showed CX-5461 treatment resulted in more cells with MK colony-forming potential **(Fig. 2G, 2H)** and increased small, medium and large colonies (**Fig. 2I, 2J**), suggesting that CX-5461 acts on undifferentiated haematopoietic stem/progenitor cells (HSPC). The data taken together indicate that CX-5461 treatment increased BM MK number and frequency, with no effect on proplatelet formation.

### CX-5461 treatment increases the LSK population and MK-biased multipotent progenitors

To further investigate the findings of increased Sca1^+^ MKs, we carried out a BM haematopoietic subpopulation analysis. A significant increase in the LSK population was identified in BM following two weeks of CX-5461 treatment (**Fig. 3A**). Using SLAM markers (CD150^+^ CD48^+^ CD135^-^) to resolve the LSK population, we observed that CX-5461 treatment led to a selective increase in the MK-primed multipotent progenitor (MPP) 2 as well as the MPP-3 population and a decrease or no change in short-term HSCs and MPP-4 populations and no change in long-term HSCs **(Fig. 3B).** Ki-67 staining showed that LT- and ST-HSCs were predominately in the G0 cell cycle phase **(Fig. 3C, 3D),** suggesting that CX-5461-treated cells were in a non-proliferative phase by day 14. Similar results were observed in MPP-2, MPP-3 and MPP-4 populations **(Fig. 3E-G).** These data suggest that CX-5461 treatment induced an MK-biased differentiation of HSPCs within the early commitment phase of haematopoiesis.

**Figure 3:**
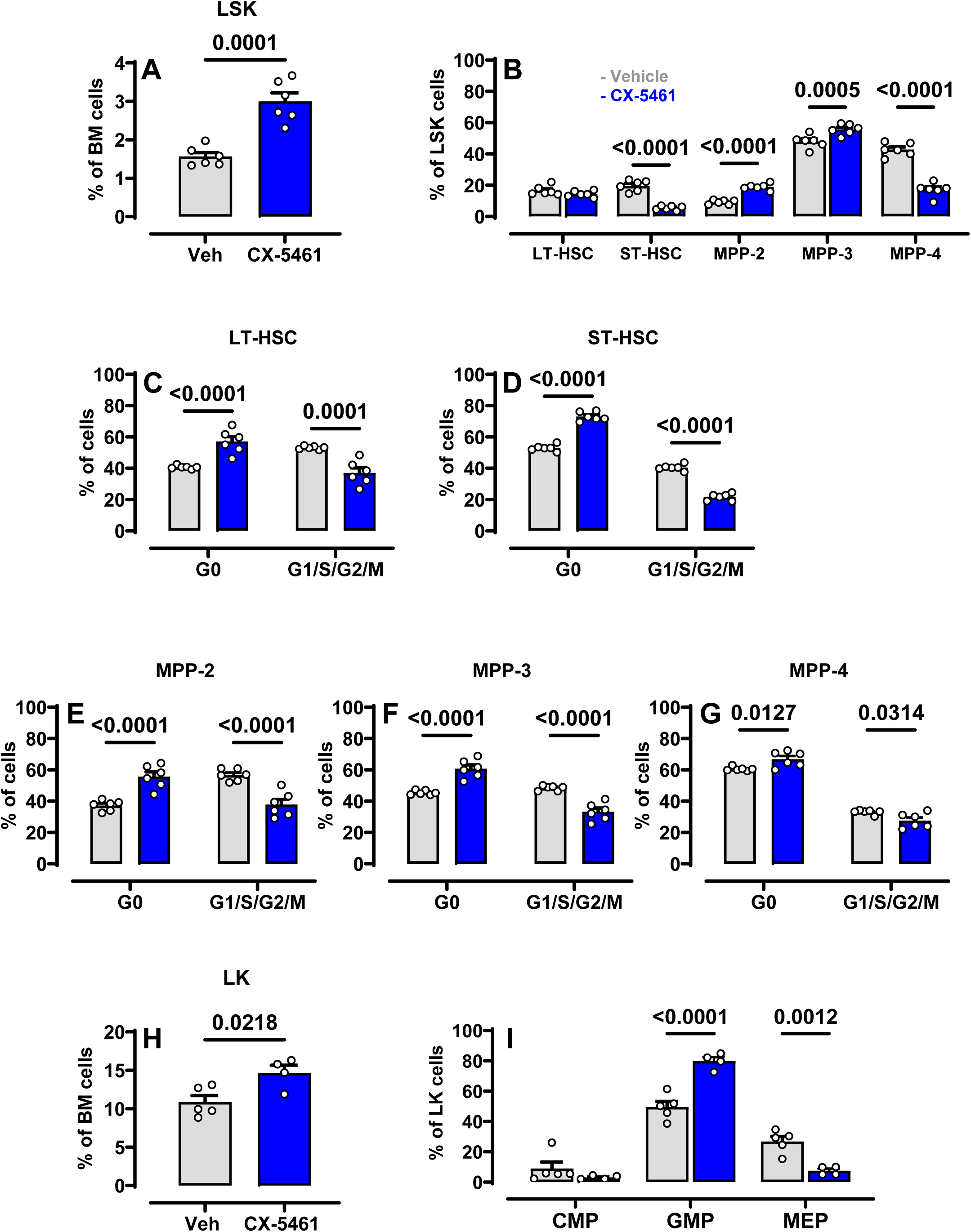
BM progenitor profile of C57BL/6 mice treated with CX-5461. Flow cytometric profiling of BM haematopoietic subpopulations in mice treated with vehicle or CX-5461 for 14 days. **(A)** Graphical representation of the LSK cell population. **(B)** LSK CD135^-^ cells were identified using SLAM markers (CD150^+^ CD48^+^ CD135^+^), LT-HSCs (LSK Flt3^-^ CD48^-^ CD150^+^), ST-HSCs (LSK Flt3^-^ CD48^-^ CD150^-^), MPP2 (LSK Flt3^-^ CD48^+^ CD150^+^), MPP3 (LSK Flt3^-^ CD48^+^ CD150^-^), MPP4 (LSK Flt3^+^) Ki-67 was used to assess the cell cycle status of each subpopulation **(C)** LT-HSC **(D)** ST-HSC **(E)** MPP-2 **(F)** MPP-3 **(G)** MPP-4. **(H)** Frequency of LK (Lin^-^Sca1^-^cKit^+^) population. LK population was analysed using CD34 and CD16/32 expression profiles to identify CMP (CD34^+^CD16/32^-^), GMP (CD34^+^CD16/32^+^) and MEP (CD34^-^CD16/32^-^). (**I)** Bar graph showing the frequency of CMPs, GMPs, and MEPs. All BM flow cytometry data and quantifications represent n= 5 to 6 mice per group. Data are presented as mean ± SEM. The significance among groups was analysed using a student’s t-test (panels A and H) or two-way ANOVA with Sidak’s multiple comparisons tests (panel B-G and I). Statistically significant numerical p-values are presented on the graphs.

CX-5461 treatment resulted in a significantly increased percentage of Lin^-^ Sca1^-^ cKit^+^ (LK) cells **(Fig. 3H).** The bipotent myeloid-erythroid progenitor (MEP) and common myeloid progenitor (CMP) pools were stable or significantly decreased **(Fig. 3I)**. Consistent with previous work demonstrating that CX-5461 treatment could induce myeloid differentiation,^15^ showed a significant increase in the granulocyte-monocyte progenitor (GMP) population in CX-5461 treated mice (**Fig. 3I**). Together, these data suggest that CX-5461 treatment prompts the activation of an MK- and myeloid-biased pathway that depletes the HSC compartment and drives the expansion of MPP-2 and MPP-3 populations to yield enhanced numbers of GMPs and MKs.

### CX-5461 treatment increased the presence of splenic MKs and MK-primed progenitors

To investigate the potential of CX-5461 treatment to alter MK abundance at sites of extramedullary thrombopoiesis, we evaluated splenic MK populations. After administering 35 mg/kg of CX-5461 for 42 days, a significant increase in spleen weight was observed **(Fig. 4A-B**). Additionally, both histology and flow cytometry confirmed a significant increase in the number of splenic MKs **(Fig. 4C-E)**. Previously splenomegaly observed in mice that were subjected to hypoxia,^18^ or infection, malignancy or inflammation,^19,20^ indicating enhanced extramedullary megakaryopoiesis. Notably, Sca1^+^ MKs were significantly increased in the spleen of CX-5461-treated mice **(Fig. 4F),** and spleen-resident progenitor analysis revealed an increased MK-primed (MPP-2) population **(Fig. 4G)** in the same mice.

**Figure 4:**
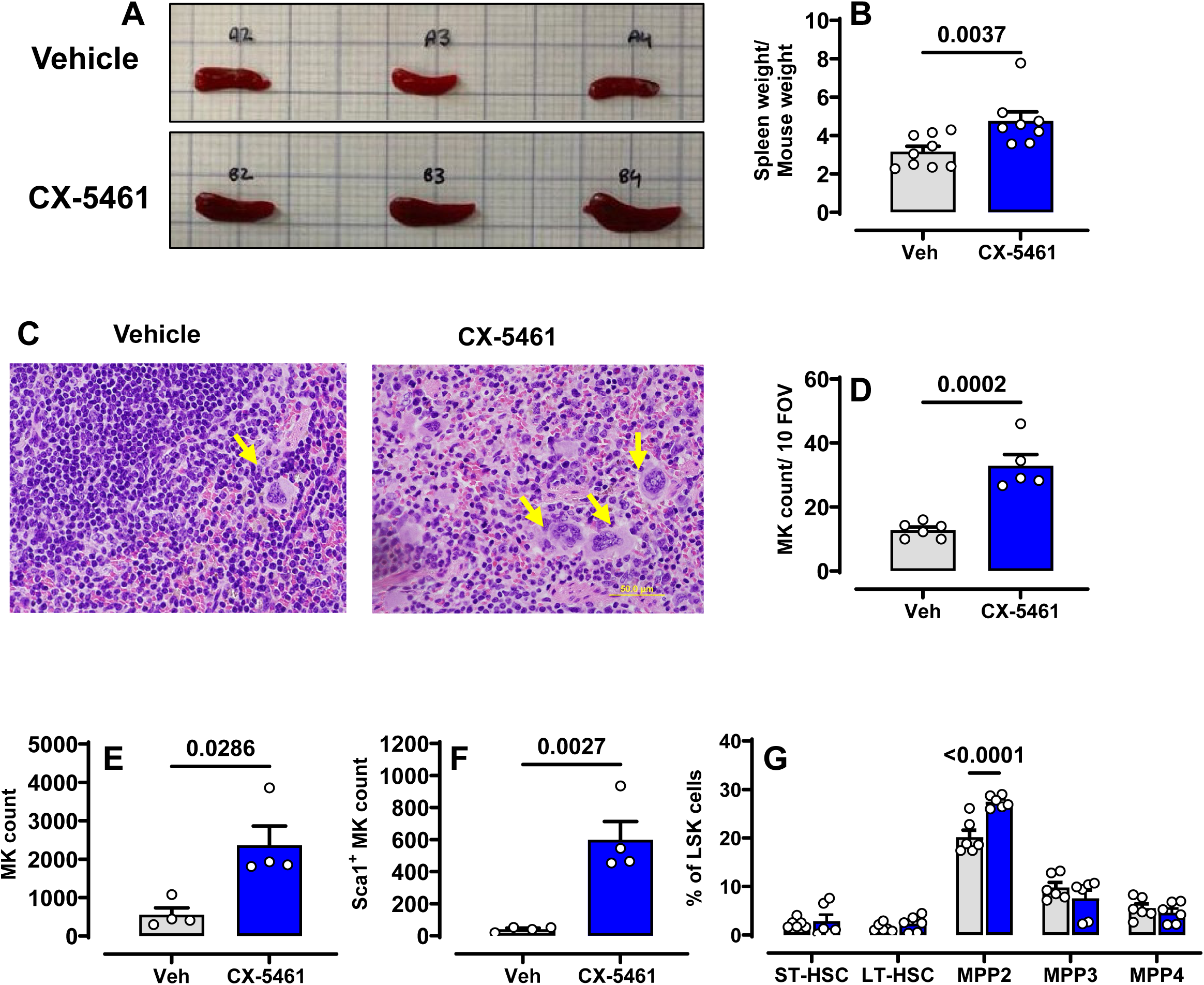
Splenic size and MK population increased in mice treated with CX-5461. C57BL/6 mice treated with 35 mg/kg CX-5461 or vehicle for 42 days. (A) Representative images of spleens at Day 42. **(B)** Spleen size and **(C)** Spleen weight normalized by body weight (mg/g) from 8 or 9 mice per group. **(D)** Representative of H&E-stained splenic sections from CX-5461 or vehicle-treated mice (n=3 mice/group). Yellow arrows indicate MKs. Scale bars, 50 µm. **(E)** Quantification of MK numbers per 10 fields of view per tissue sample from n = 4 mice/group. Analysis was conducted by an operator who was blinded to the treatments. **(F)** Splenic MK counts (G) Sca1^+^ cells within the splenic MK population. **(H)** Quantification of ST-HSCs, LT-HSC, MPP2, MPP3, MPP4 was performed following 2 weeks of CX-5461 or vehicle treatment. All flow cytometry data were derived from 4 or 5 mice per treatment group. Data are presented as mean ± SEM. The significance among groups was analysed by student’s t-test (panels E, and G) or Mann Whittney’s test (panels B, C and F) and two-way ANOVA with Sidak’s multiple comparisons (panel H). Statistically significant numerical p-values are presented on the graphs.

### The thrombopoietic effect of CX-5461 is independent of TPO and c-mpl signalling

TPO is the primary regulator of MK/platelet biogenesis and is involved in the self-renewal and maintenance of HSCs.^21^ Hepatocyte Tpo mRNA and plasma TPO levels were assessed by RT-PCR and ELISA, respectively. TPO mRNA and TPO protein levels were comparable between CX-5461- and vehicle-treated groups **(ED. 4),** suggesting that CX-5461 treatment did not accelerate the production of TPO.

To examine the role of TPO/c-mpl signalling, c-mpl-deficient mice (c-mpl^-/-^) were treated with vehicle or CX-5461 for up to 28 days. As reported by others,^4^ untreated c-mpl^-/-^ mice bore approximately 10% of an average murine platelet count (**Fig. 5A**). As observed in wild-type (WT) mice, CX-5461 c-mpl^-/-^ mice treatment resulted in rapidly increased circulating platelet numbers that were reversible with treatment cessation and reduced or unchanged levels of RBC and lymphocytes **(Fig. 5B, C).** While the platelet counts consistently remained beneath platelet levels typically observed in WT mice (**Fig. 1A**), c-mpl^-/-^ mice displayed a 3.5-fold increase in platelet levels after 28 days of exposure to CX-5461, indicating a robust response to CX-5461 treatment in thrombocytopenic mice. The increased platelet count was accompanied by an increase in the immature platelet fraction **(Fig. 5D).** Platelets produced under these conditions displayed normal levels of platelet receptors, no evidence of enhanced platelet activation and normal responses to collagen **(Fig. 5E, 5F).**

**Figure 5:**
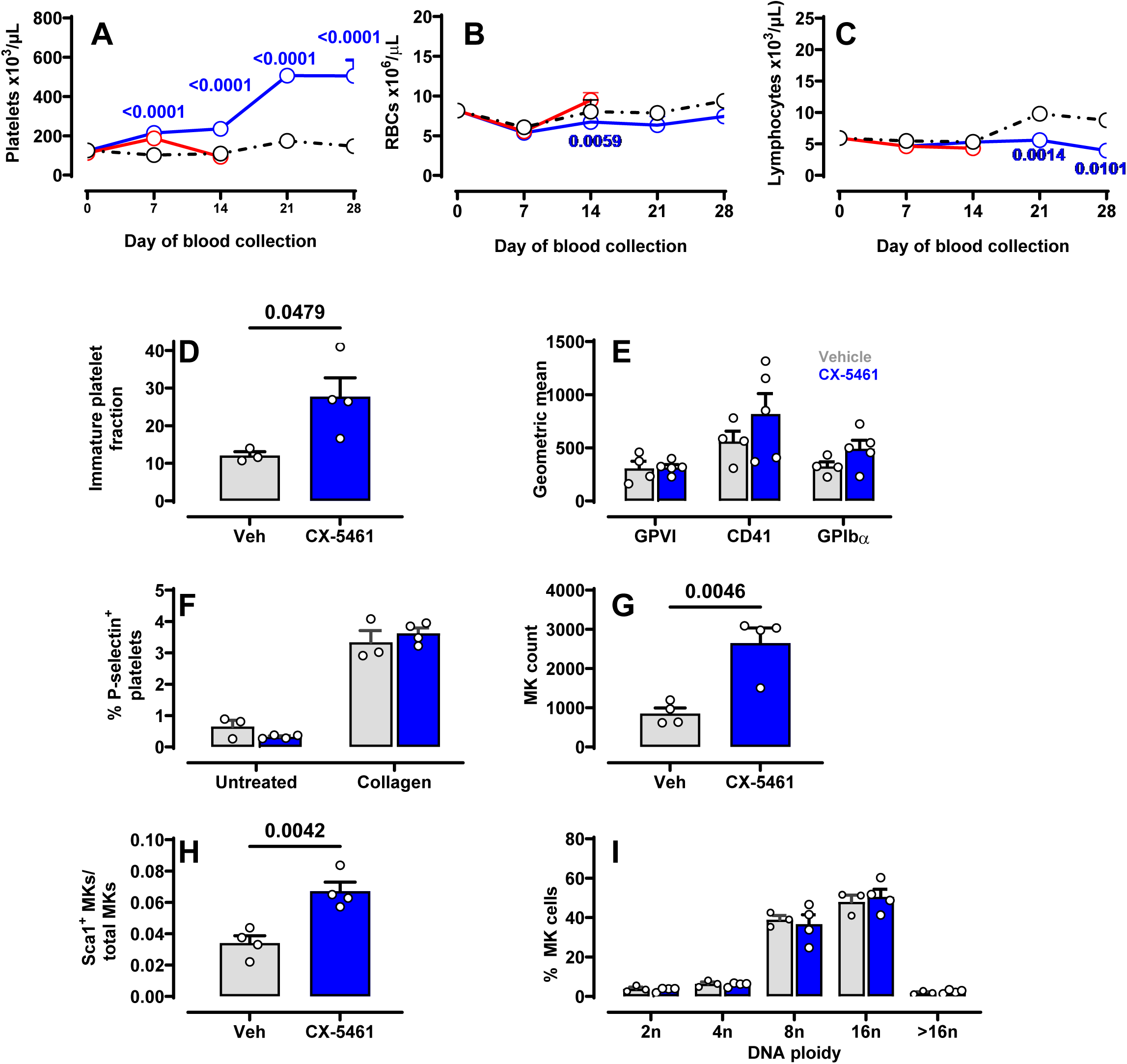
CX-5461 treatment increases platelets and MK numbers in *c-mpl* ^-/-^ mice. C57BL/6 *c-mpl^-/-^* mice (n=3-5 per group) were treated with 35 mg/kg CX5461 for 1 week (blue line) or 4 weeks (red) or vehicle (black line). Blood samples were collected on days 0, 7, 14, 21, and 28, and **(A)** platelets, **(B)** RBCs, and **(C)** lymphocytes were quantified. At day 28 peripheral blood and BM samples were collected from the vehicle and CX-5461 treated mice for the following quantifications through flow cytometry. **(D)** Quantification of immature platelet fraction based on thiazole orange staining. **(E)** Surface levels of receptors GPVI, CD41 (integrin αIIb), and GPIbα on circulating platelets. **(F)** P-selectin levels on resting or 10 µg/mL collagen-activated platelets. (**G)** BM MK numbers and **(H)** frequency of CD41^+^ Sca1^+^ MKs within each MK population quantified from 10^6^ BM cells. **(I)** DNA ploidy of MKs was quantified using PI staining in a flow cytometer (n=4-5). Data are represented as mean ± SEM. The significance among groups was analysed using student’s t-test (panels D, G, and H) or Two-way ANOVA with Sidak’s (panels E, F, and I) or Tukey’s multiple comparisons (panels B and C) or Dunnett’s multiple comparisons (panel A). Statistically significant numerical p-values are presented on the graphs.

Like WT mice, c-mpl^-/-^ mice treated with CX-5461 for 28 days significantly increased MK numbers in BM compared to vehicle-treated mice **(Fig. 5G)**, with increased Sca1^+^ MKs **(Fig. 5H)** but no changes in MK ploidy (**Fig. 5I**). These data suggest that CX-5461 treatment increases the numbers of platelets, MKs and Sca1^+^ MKs, but not MK maturation and this effect was independent of TPO/c-mpl signalling.

Large increases in specific cytokines such as interleukin (IL)-6 may lead to enhanced megakaryopoiesis.^22^ We assessed levels of 23 cytokines in serum and BM samples from vehicle-or CX-5461-treated WT and c-mpl^-/-^ mice **(STables 3, 4)**. CX-5461 treatment in WT mice was associated with a reduction in serum levels of C-C motif chemokine ligand 5 (CCL5) and an increase in CCL11, with no significant alterations observed in other cytokines. In corresponding BM supernatants, increases in interleukin (IL)-1α, CCL11, CCL5, keratinocyte chemoattractant, and monocyte chemoattractant protein (MCP)-1 were detected (**STable 3)**. In c-mpl^-/-^ mice, elevated levels of serum keratinocyte chemoattractant and increased BM supernatant concentrations of MCP-1 **(STable 4)**. Whilst some treatment-induced changes were observed, the fold-changes in cytokines were modest and within physiological ranges. We concluded that the upregulation of inflammatory cytokines did not significantly contribute to CX-5461-induced thrombopoiesis.

### CX-5461 treatment promotes platelet rescue following carboplatin-induced thrombocytopenia

Thrombocytopenia is a common and challenging complication of cancer therapies, often limiting treatment intensity and increasing the risk of bleeding, which can compromise patient outcomes.^23^ We explored the efficiency of CX-5461 as an adjunct therapy to rescue chemotherapy-induced thrombocytopenia. Mice were treated with chemotherapeutic reagent carboplatin alone or together with 15 mg/kg CX-5461. Carboplatin induced reductions in platelet numbers as expected. However, CX-5461-treated mice exhibited a reduced depth of thrombocytopenia at the nadir (day 8) and a more rapid rate and extent of platelet count rebound relative to carboplatin treatment alone **(Fig. 6A)**. Analysis of the BM showed comparable levels of BM MKs **(Fig. 6B)** in carboplatin-treated mice ± CX-5461 possibly owing to lower doses of the CX-5461 or the limited duration of treatment and observation. While LT-HSCs showed no significant differences **(Fig. 6C)**, a marked reduction in ST-HSCS frequency was observed (**Fig. 6D**), alongside a significant increase in MPP-2 and MPP-3 populations **(Fig. 6E, 6F)** in CX–5461–treatment with carboplatin, compared to carboplatin-only controls. Carboplatin treatment severely ablated the MPP-4 subpopulation **(Fig. 6G)**. Under these experimental conditions, CX-5461 treatment effectively rescues platelet count in chemotherapy-exposed thrombocytopenic mice, most likely by driving differentiation of ST-HSCs towards MPP-2 and MPP-3 subpopulations.

**Figure 6:**
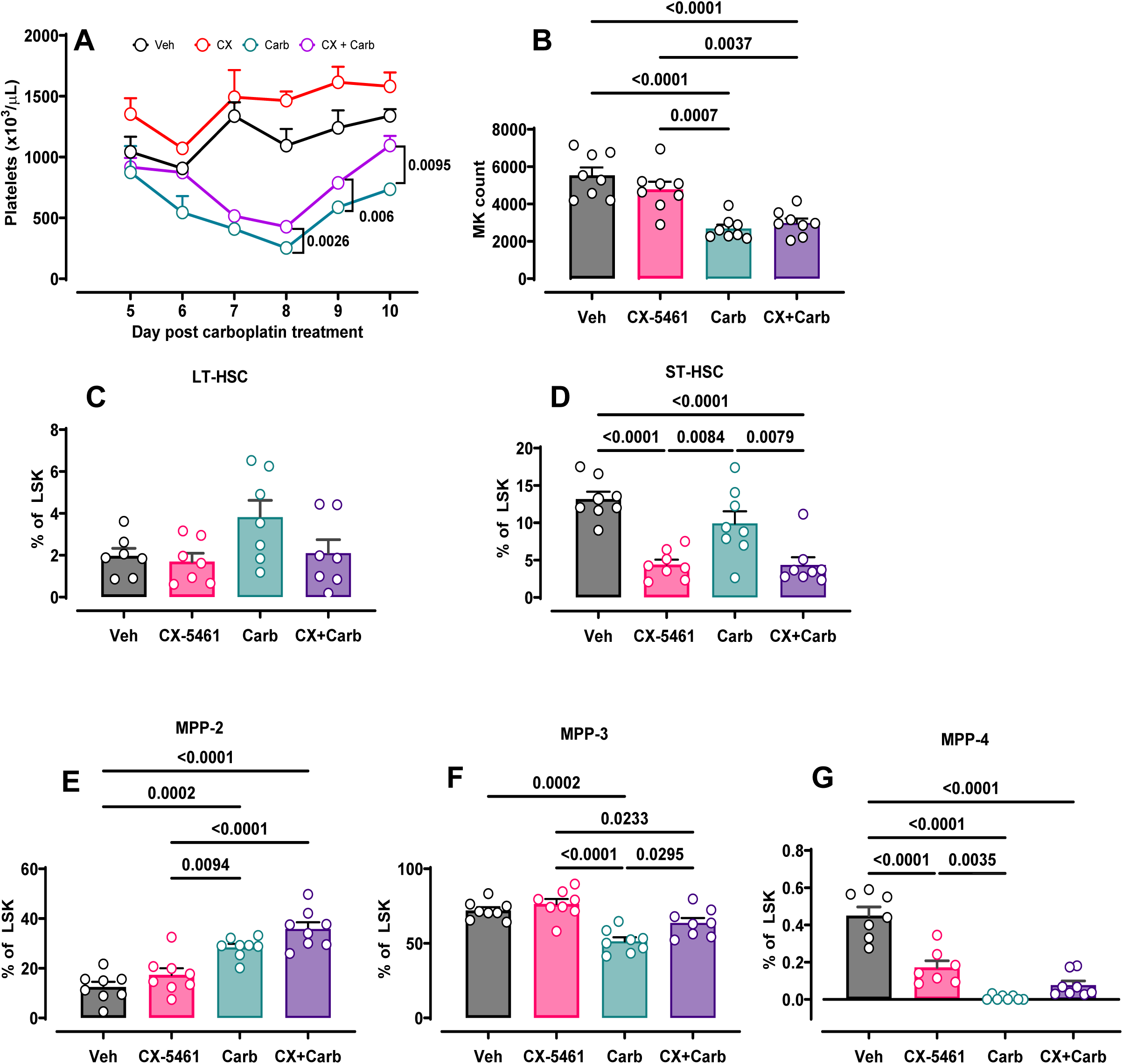
CX-5461 treatment accelerates platelet rescue following chemotherapy-induced thrombocytopenia. Mice receiving 15 mg/kg CX-5461 (3 doses/week) were I.P. injected with 80 mg/kg carboplatin or vehicle for 10 days. Mouse tail vein blood counts were collected at different time points, and BM cell phenotyping was performed 10 days after carboplatin treatment. **(A)** Line graph showing temporal changes in platelet count following carboplatin treatment. Flow cytometry quantification of mouse BM **(B)** MKs **(C)** LT-HSC **(D)** ST-HSC **(E)** MPP-2 **(F)** MPP-3 **(G)** MPP-4 (n=4-8). Data are represented as mean ± SEM. The significance among groups was analysed using two-way ANOVA with Sidak’s multiple comparisons (panel A) or One-way ANOVA with Tukey’s multiple comparisons test (panels C-G). Statistically significant numerical p-values are presented on the graphs.

### CX-5461 treatment decreased MK cell proliferation and maturation in vitro

We evaluated the direct effects of CX-5461 on MK precursor proliferation and differentiation using both an immortalised human MK cell line (imMKCL) and cultured mouse BM cells. Under cell proliferative conditions (+ doxycycline), imMKCL cells were treated with 50-300 nM CX-5461. CX-5461 treatment resulted in a dose-dependent inhibition of imMKCL proliferation (**ED. 5A**). Under conditions of maturation (no doxycycline), CX-5461 did not affect the expression of venus-tagged β-tubulin, a MK maturation reporter and GPIbα surface expression, suggesting no impact of CX-5461 on MK cell maturation (**EDs. 5B, 5C**). Findings suggest that CX-5461 treatment negatively impacted MK proliferation but not maturation.

### CX-5461 treatment enhances the differentiation of MkP to MKs

To elucidate the cellular and molecular mechanisms driving CX-5461-induced MK lineage-biased haematopoiesis, we performed high-throughput single-cell RNA sequencing (scRNA-seq) of LSK cells isolated from the BM of mice treated with CX-5461 or vehicle for 14 days, performed in four biological replicates. Using UMAP for dimensionality reduction and Leiden algorithm for clustering, we identified 21 distinct cell populations, which were annotated based on differentially expressed genes and reference marker genes from the CellKb database (**ED. 6A** and **STable 5**). Among these, two specific megakaryocytic clusters, megakaryocyte progenitors (MkPs) and mature MKs, were identified. The relative proportions of these clusters were similar between treatment groups (**ED. 6B**).

To assess lineage dynamics, we applied RNA velocity analysis on the LSK population, a method that infers future transcriptional states of individual cells by analysing the ratio of unspliced to spliced mRNA. This analysis revealed CX-5461 treatment induced a directional shift in transcriptional kinetics, promoting a transition from MkPs toward MKs (**ED. 6C**). Further lineage inference using velocity pseudotime analysis and partition-based graph abstraction (PAGA) analysis, using HSCs as the root, demonstrated a strong propensity for MkP to differentiate into MK cells in response to CX-5461 treatment (**Fig. 7A, ED. 6C**). These results align with the increased MK numbers and MK-biased progenitor populations observed, supporting that CX-5461 enhances the differentiation process from MkPs to MKs.

**Figure 7:**
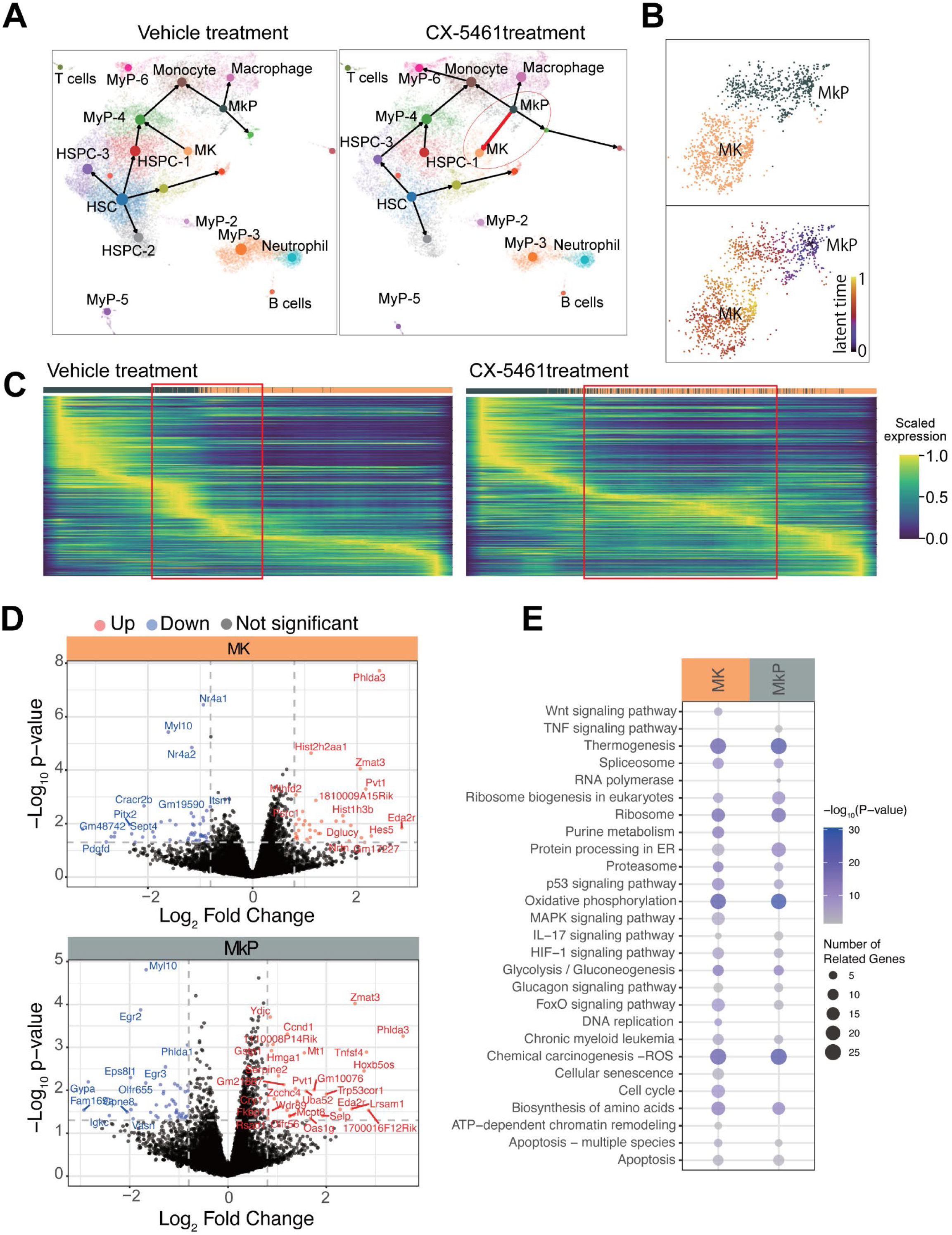
scRNA sequencing CX-5461 treatment promotes the differentiation of MKPs to MKs. C57BL/6 mice were treated with 35 mg/kg of CX-5461 or vehicle for 14 days and BM LSKs were sorted and processed for scRNA sequencing. **(A)** UMAP plots of vehicle control (left) and CX-5461-treated (right) samples, with lineage tracing based on pseudo-time analysis indicated by arrows suggesting putative cell transitions. **(B)** UMAP plots focusing on MK and MKP clusters (top), and corresponding visualisation of latent time dynamics within these clusters (bottom). **(C)** Heatmaps showing the scaled expression levels of the top 300 genes identified as ’driver genes’ ranked by latent time. These genes exhibit pronounced dynamic behaviour across the dataset and are systematically selected based on their high fit likelihoods in the dynamic model, ranked by latent time in both vehicle (left) and CX-5461-treated (right) samples. Expression levels are convolved over 100 cells to enhance visual clarity. **(D)** Volcano plots showing significantly differentially expressed genes in MkP and MK clusters between vehicle and CX-5461 treatments. Gene significantly upregulated and downregulated following CX-5461 treatment shown in red and blue, respectively. **(E)** Pathway enrichment analysis identifies pathways significantly enriched in differentially expressed genes in MkP and MK clusters following CX-5461 treatment.

We next focused on the MK clusters and used a dynamical model to examine changes in latent time, a measure of a cell’s progression through a biological process **(Fig. 7B)**. Ranking the top 300 genes by latent time revealed distinct expression dynamics: in vehicle-treated mice, MK cells exhibited gradual gene expression changes along the latent trajectory, consistent with steady-state differentiation **(Fig. 7C)**. In contrast, CX-5461-treated cells showed a marked shift toward later latent stages, indicating accelerated MkP-to-MK maturation. This acceleration was corroborated by an elevated presence of progenitor cells at latent time points corresponding to mature MKs, with heatmaps highlighting a more pronounced transition from early to late stages in the CX-5461-treated group (red rectangles, **Fig. 7C**). This finding is consistent with the rapid elevation in platelet counts and enhanced megakaryopoiesis observed in the BM of CX-5461-treated mice.

Finally, differential gene expression analysis of the MK and MkP clusters revealed significant gene expression changes between CX-5461- and vehicle-treated samples (**Fig. 7D**). Notably, CX-5461 treatment resulted in the enrichment of key pathways, including oxidative phosphorylation, p53 signaling, and ribosome biogenesis (**Fig. 7E**), primarily associated with upregulated differentially expressed genes **(STable 6)**.

## Discussion

In this study, we report the discovery of a novel thrombopoietic effect of the RNA Pol I inhibitor CX-5461, a compound originally developed to target ribosome biogenesis in cancer. Our data reveal that CX-5461 not only exerts potent anti-tumour activity but also significantly enhances megakaryopoiesis and increases platelet counts in both humans and mice, through a mechanism that is independent of thrombopoietin (TPO) or c-mpl receptor signalling.

This observation was first made in patients enrolled in a phase I dose-escalation trial, where over half of the participants exhibited an increase in peripheral platelet counts following a single administration of CX-5461. Importantly, this increase occurred in the absence of systemic inflammation or evidence of platelet activation, suggesting that the effect was a *bona fide* physiological enhancement of platelet production. We confirmed and extended this finding in preclinical mouse models, where CX-5461 treatment elicited a rapid, sustained, and reversible increase in circulating platelets. Platelet function, surface marker expression, and half-life were preserved, indicating that the drug promotes production of functional platelets without compromising quality or turnover.

Mechanistically, we show that CX-5461 enhances the number of BM MK, particularly the Sca1⁺ MK subset, and expands megakaryocyte-biased multipotent progenitors (MPP-2 and MPP-3). These changes accompanied an increased frequency of LSK (Lin⁻ Sca1⁺ cKit⁺) cells and an augmented MK differentiation trajectory revealed by single-cell RNA-seq and RNA velocity analysis. Strikingly, these effects were also observed in c-mpl⁻/⁻ mice, establishing that CX-5461 acts independently of the TPO receptor. Together, our data support a model in which CX-5461 modulates HSC and HSPC fate, promoting MK lineage bias through TPO-independent pathways.

Our findings extend a growing body of literature demonstrating that inhibition of Pol I transcription can trigger differentiation in multiple biological contexts. Hein and colleagues demonstrated that Pol I inhibition disrupts self-renewal in acute myeloid leukaemia by promoting differentiation and depleting leukaemia-initiating cells.^15^ Hayashi and colleagues showed that reduced rDNA transcription induces differentiation of embryonic stem cells,^24^ and Zhang and colleagues found similar lineage-defining effects in *Drosophila* stem cell systems.^25^ Our work builds on these foundational studies, revealing that Pol I inhibition also primes haematopoietic progenitors towards megakaryopoietic fate *in vivo*.

In the canonical pathway of haematopoiesis, HSCs give rise to CMP and CLP populations from which granulocytes and the MEP population, or lymphocyte and natural killer cell populations arise, respectively. MEPs may also arise directly from HSCs^26^ and using single-cell genomics and lineage tracing and computational tools, lineage-biased haematopoietic pathways are now under intensive investigation.^27^ MK-biased HSCs are characterized by expression of the αIIb integrin subunit (CD41), von Willebrand Factor (VWF) and Sca-1 as well as their location in the BM^28^ and their response to TPO receptor engagement.^29^ These MK-primed HSCs differentiate directly into MKs, bypassing intermediate progenitor populations, and produce platelets under healthy conditions.^30^ This subset proliferates in response to the addition of TPO to HSC cultures *in vitro,*^31^ toll-like receptor (TLR) activation^21^ or type I interferon^32^ and is restrained by selective chromatin compaction within regulatory regions controlling platelet genes.^33^ Here BM from CX-5461-treated mice showed elevated MK, MPP-2, and myeloid subpopulations and an increased propensity towards megakaryopoiesis with high MK CFUs. This evidence aligns closely with the non-canonical pathway of MK/platelet-restricted HSC fate commitment.^28,34^ Discontinuation of CX-5461 treatment resulted in the normalization of platelet numbers and restoration of erythrocyte and leukocyte counts to normal levels consistent with resumption of normal haematopoiesis. Furthermore, single-cell transcriptomic analysis identified that CX-5461 treatment augmented a differentiation trajectory of MKP towards MKs. Overall, this work suggests that CX-5461 treatment potentiates the MK/platelet lineage biased haematopoietic output.

Megakaryopoiesis and thrombopoiesis predominantly occur in BM as part of haematopoiesis. rRNA gene transcription into pre-rRNA (47S rRNA) by RNA Pol I influences haematopoiesis and plays a critical role in stem cell fate decisions and lineage specification.^35^ MKs of high ploidy exhibit enhanced ribosome biogenesis and protein translation,^36^ and we noted changes to MK numbers but not MK ploidy. Our work identified increases in numbers of LSK cells in CX-5461-treated mice that were predominantly in a non-proliferative (G0) phase. Despite HSCs exhibiting a reduced proliferative capacity compared to actively dividing MPPs, these cells reportedly maintain similar levels of rDNA transcription.^37^ Most HSCs contain established megakaryotic gene profiles, control of MK maturation is under the influence of transcription factor RUNX1,^38^ and extensive *de novo* DNA methylation is necessary to induce megakaryopoiesis.^39^ Future work will assess changes in haematopoietic-biased transcription factor profiles and the DNA methylomes in HSC from mice treated with CX-5461.

Transcriptomic data from our single-cell RNA-seq analysis indicated that genes within p53 signalling pathways were primarily upregulated under conditions where enhanced differentiation of MkP to MKs was observed. While we have not experimentally proven a causal relationship between these two observations, it mirrors recent findings showing elevated p53 signalling and unfolded protein response (UPR) in VWF^+^ platelet/MK-biased HSCs.^34^ CX-5461 triggers nucleolar stress that induces DNA damage and cell cycle arrest at G2, two factors that are known to influence rates and extent of MK differentiation.^40^ This evidence suggests that p53 may plays a fundamental role in directing haematopoietic differentiation towards the MK lineage in response to cellular stress. Notably, the observed p53 upregulation supports a model where nucleolar stress shifts HSC fate toward the MK lineage. Furthermore, p53’s role could intersect with other stress pathways, such as the UPR, potentially amplifying MK maturation under CX-5461 treatment, as hinted by the accelerated latent time trajectories in our transcriptomic data. These findings position p53 as a central mediator in CX-5461’s TPO-independent thrombopoietic effects, consistent with its action in c-mpl-deficient mice. Targeting p53-related pathways could enhance therapeutic strategies for thrombocytopenia, particularly in chemotherapy-induced settings where MK differentiation is critical.

Thrombocytopenia can be caused by factors relating to viremia, drug- or therapeutic exposure as well as patient-specific features such as autoantibodies.^41^ TPO receptor agonists are the major approved drugs used to increase platelet counts,^42^ in acute and chronic thrombocytopenia settings when thrombocytopenia poses a bleeding risk or limits treatment options. TPO mimetics are effective in immune thrombocytopenia but show inconsistent results in conditions like myelodysplastic syndromes or aplastic anaemia. Some patients exhibit primary or secondary resistance to these agents. To overcome these limitations, novel drugs (stand-alone or in combination) that act via TPO receptor-independent pathways are needed. Our work shows CX-5461 could be explored for thrombocytopenia treatments. Here, carboplatin-mediated reduction in circulating platelet numbers was offset by treatment with a low dose of CX-5461 suggesting utility of ribosome biogenesis inhibitors in chemotherapy-induced thrombocytopenia settings. CX-5461 may not be the ideal candidate due to drug-induced mutagenesis effects^43^ however we have data showing that other Pol I inhibitors stimulated thrombopoiesis. Excitingly, new generation specific Pol I inhibitors with improved pharmacokinetic and specificity profiles are emerging,^11,44^ and await testing in the haematopoiesis setting.

Limitations of this study include data on the effect of RNA Pol I inhibitors on haematopoiesis in mice, with limited information on these inhibitors in humans. The presented human data involved patients from clinical trials who had terminal diseases, and samples were limited to those available from collaborators at other institutes and not available for all study participants. This work assumes a causal relationship between MPP-2, MKs, Sca1^+^ MKs and peripheral platelet numbers however this inference is accepted practice in existing literature. BM transplantation and limited dilution experiments will bolster evidence that the thrombopoietic effect manifests at the level of HSCs and that an HSC-MK-biased population is expanded by CX-5461 exposure.

The novel thrombopoietic effects of Pol I inhibition advance our understanding of the complex interplay between the rate and regulation of ribosome biogenesis in haematopoiesis. While disruption of ribosome biogenesis, such as that observed in ribosomopathies, is frequently associated with cytopenias, our work demonstrates an unappreciated capacity to promote myeloid and megakaryocytic differentiation. CX-5461 is a novel pharmacological agent capable of inducing TPO-independent MK-biased thrombopoiesis, thus a valuable tool for dissecting the intricate mechanisms that govern platelet production and potentially a prototype molecule for new thrombopoiesis therapeutics.

## Materials and Methods

Additional methods are detailed in Supplementary Methods.

### CX-5461 human trial samples

Blood samples from haematology patients were obtained after provision of written informed consent under Protocol No. PMCC 12/79 from the Peter MacCallum Cancer Centre as part of a phase I clinical trial evaluating CX-5461 (Australia and New Zealand Clinical Trials Registry, #12613001061729).^9^ A full blood cell analysis was performed, and double-spun plasma was isolated and stored at −80°C.

### Animal experiments

All animal experiment procedures were approved by the Australian National University (ANU) Animal Experimentation Ethics Committee (A2018/48; A2021/42). C57BL/6 and C57BL/6 c-mpl*^-/-^* (tm1Wsa)^45^ mice of mixed gender, aged 9 to 18 weeks, were bred in-house at ANU and housed within physical containment (PC)2 facilities.

### CX-5461 and BMH-21 treatment in mice and blood collection and analysis

Groups of five to eight mice were treated with 10-35 mg/kg CX-5461, 75 mg/kg BMH-21 or an equivalent volume of 50 mM Na_2_PO_4_ buffer, pH 4.5 (vehicle) via oral gavage routinely 3 times/week. Murine tail vein blood was collected as described previously.^46^ Blood cell quantitation was performed using an automated blood analyzer (Advia 2120, Siemens Australia). Diluted whole blood was incubated with agonists (platelet activation studies), thiazole orange (immature platelet fraction analysis) or fluorescently labelled antibodies. Platelet surface proteins were analyzed on an LSRII flow cytometer (Becton Dickinson).

### Enumeration of spleen and BM MKs

Homogenized spleen and BM samples harvested from femora and tibiae of euthanized mice were processed by centrifugation^47^ and subjected to hypotonic lysis by resuspension in a solution of 17 mM Tris-HCl, pH 7.6 containing 144 mM NH_4_Cl for 5 minutes at 4°C followed by centrifugation and resuspension in CATCH buffer (in mmol L-1: 5.3 KCl, 0.44 KH_2_PO_4_, 137 NaCl, 4.17 NaHCO_3_, 0.338 Na_2_HPO_4_, 5.56 Glucose, 12.9 sodium citrate, 1.0 adenosine, 2.0 theophylline, 3% FBS [v/v], 3% BSA [w/v]). Cell suspensions were mixed with antibodies against CD41, stem cell antigen (Sca)-1 (**Supp. Table 4**) and propidium iodide (PI), and the frequency of MKs were analyzed by flow cytometry. For MK ploidy assays, BM isolates were resuspended in 0.1% sodium citrate containing 50 µg/mL PI and 50 µg/mL RNase for 30 mins at 4°C and analyzed by flow cytometry where MKs were gated on forward and side scatter properties and CD41 positive staining. MK DNA content (ploidy) was measured by PI staining, and maturation was assessed based on CD42b expression. To visualize and enumerate MKs histologically, femurs and spleens were fixed in 10% formalin, then sectioned longitudinally, and stained with haematoxylin and eosin (H&E). Using light microscopy (Olympus XI7, 100X magnification), an operator blinded to sample treatments counted the number of MKs per field of view across 28 fields/section.

### Evaluation of BM haematopoietic progenitor and stem cells

BM isolates underwent RBC lysis followed by lineage depletion using FITC-labelled antibodies and anti-FITC MACS beads and then passed through LS columns (Miltenyi Biotec, Australia) as per the manufacturer’s protocol. Cell suspensions were subsequently stained with antibodies against c-Kit, Sca1, CD34, CD135/Flt3, CD48, CD150, CD41, CD127 and CD16/32 for 30 minutes in the dark at 4°C followed by live-dead Aqua staining. Cells were centrifuged, resuspended and then immunosorted (FACS Aria II or FACSFusion, Becton Dickinson). HSCs were gated as a lineage-negative (Lin^-^) Sca1^+^cKit^+^ (LSK) population. MPPs were gated based on SLAM markers (CD150^+^ CD48^+^ CD135^+^) as described.^48^ The common myeloid progenitor (CMP), granulocyte-monocyte progenitor (GMP), and MK-erythroid progenitor (MEP) were gated as described.^47^ ^47^

### Platelet half-life *in vivo*

One-week post vehicle or 35 mg/kg CX-5461 treatment, mice were injected intravenously with 0.15 µg/g of Dylight-488-conjugated anti-CD42c antibody recognising platelet-specific receptor GPIbβ to label all platelets. Blood samples were periodically collected and mixed with an anti-CD41 antibody. Platelets were gated on CD41 antibody positivity and analysed for DyLight-488 anti-CD42c binding.

### Statistical analysis and software utilised

Statistical analyses were performed using GraphPad Prism v8.0c (GraphPad Software, California, USA). Data were evaluated for normal distribution using the Shapiro-Wilk test, and appropriate parametric or non-parametric statistical analyses on raw data were applied as described in Figure legends. Data are represented as mean±SEM. Microscopic images were analysed using Image J Fiji (NIH, Maryland, USA). Flow cytometry data were analysed using FlowJo v10.2 (FlowJo, LLC Oregon, USA).

## Acknowledgements

We thank Mr M. Devoy and Dr H. Vohra of the Cytometry Histology and Spatial Microscopy Facility and Dr Maxim Nekrasov from the Biomedical Resource Facility at the John Curtin School of Medical Research. This work was funded by the National Health and Medical Research Council of Australia, the Cancer Council ACT, and the Bootes Family, Canberra, Australia.

## Author contribution

VB, AK, and EEG wrote the original draft; VB, AK, JIH, RF, NH, KHM, SN, SAA, SAB and KMH carried out vital experiments; LL and JW provided analyses of scRNAseq data; SMM, SJH, KE provided key reagents; AK, RF, RKA, PYIC, KMH, RDH and EEG acquired funding and directed the research. All authors reviewed and interpreted the data. During the writing of this manuscript, KMH died. All other authors reviewed and approved the final manuscript.

## EXTENDED DATA FIGURES

**Extended Data 1:**
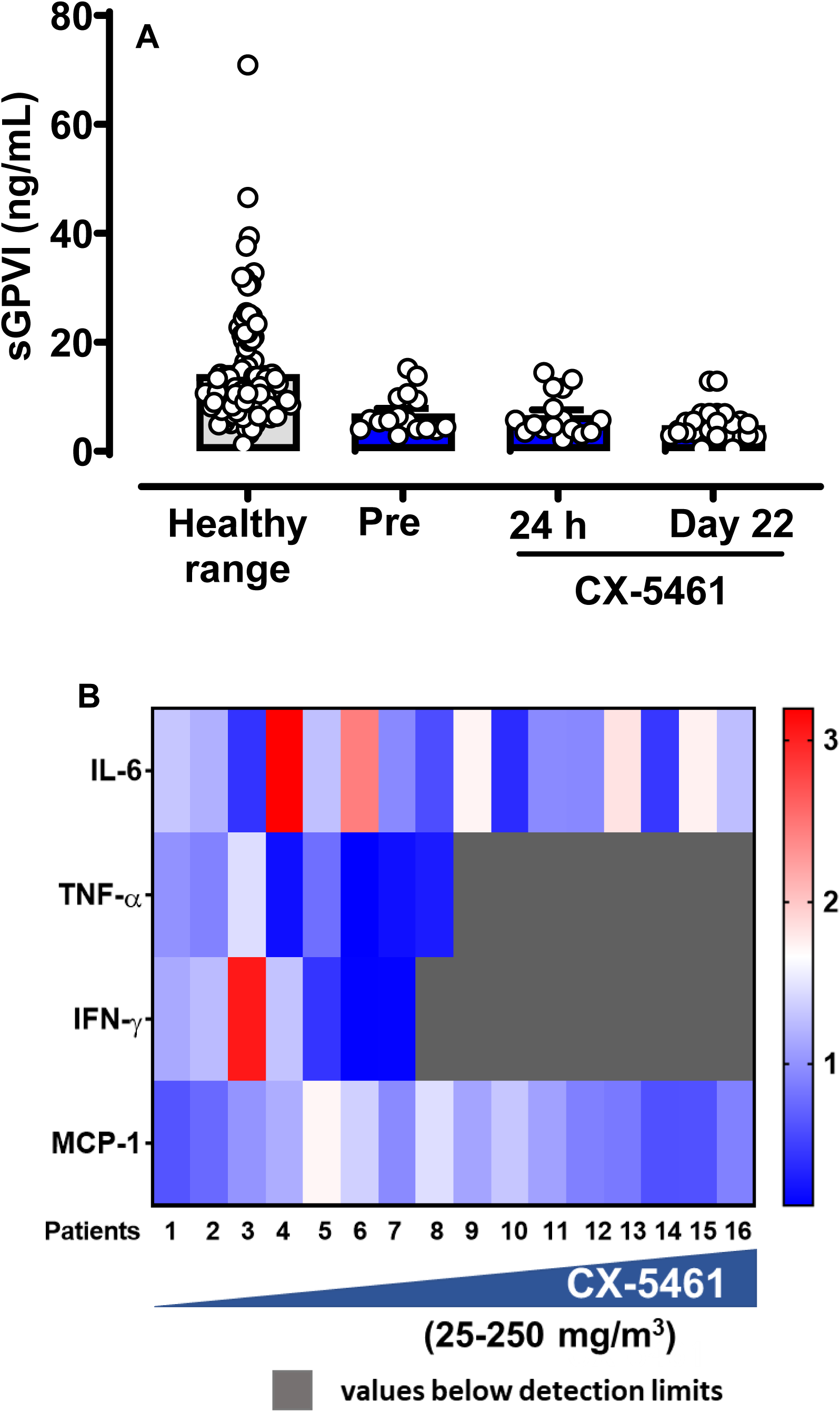
Patients treated with CX-5461 show low levels of sGPVI and no significant change in inflammatory markers in their plasma. **(A)** Plasma sGPVI levels in 16 patients with advanced haematological malignancy were measured in samples isolated prior to treatment (pre), or 24 h or 22 days post-drug treatment by ELISA. Data are presented as mean ± SEM. The patients received single doses ranging from 25-250 mg/m^2^ CX-5461. **(B)** Plasma isolated from patients’ blood 24 hours after CX-5461 treatment, were assessed for cytokines IL-6, TNF-α, IFN-γ, and MCP-1 using a multiplexed ELISA as described in methods. Levels are shown as heat maps. Statistical differences were analysed using the Kruskal Wallis test with Dunn’s multiple comparisons between samples collected at three time points. Statistically significant numerical p-values are presented on the graphs.

**Extended Data 2:**
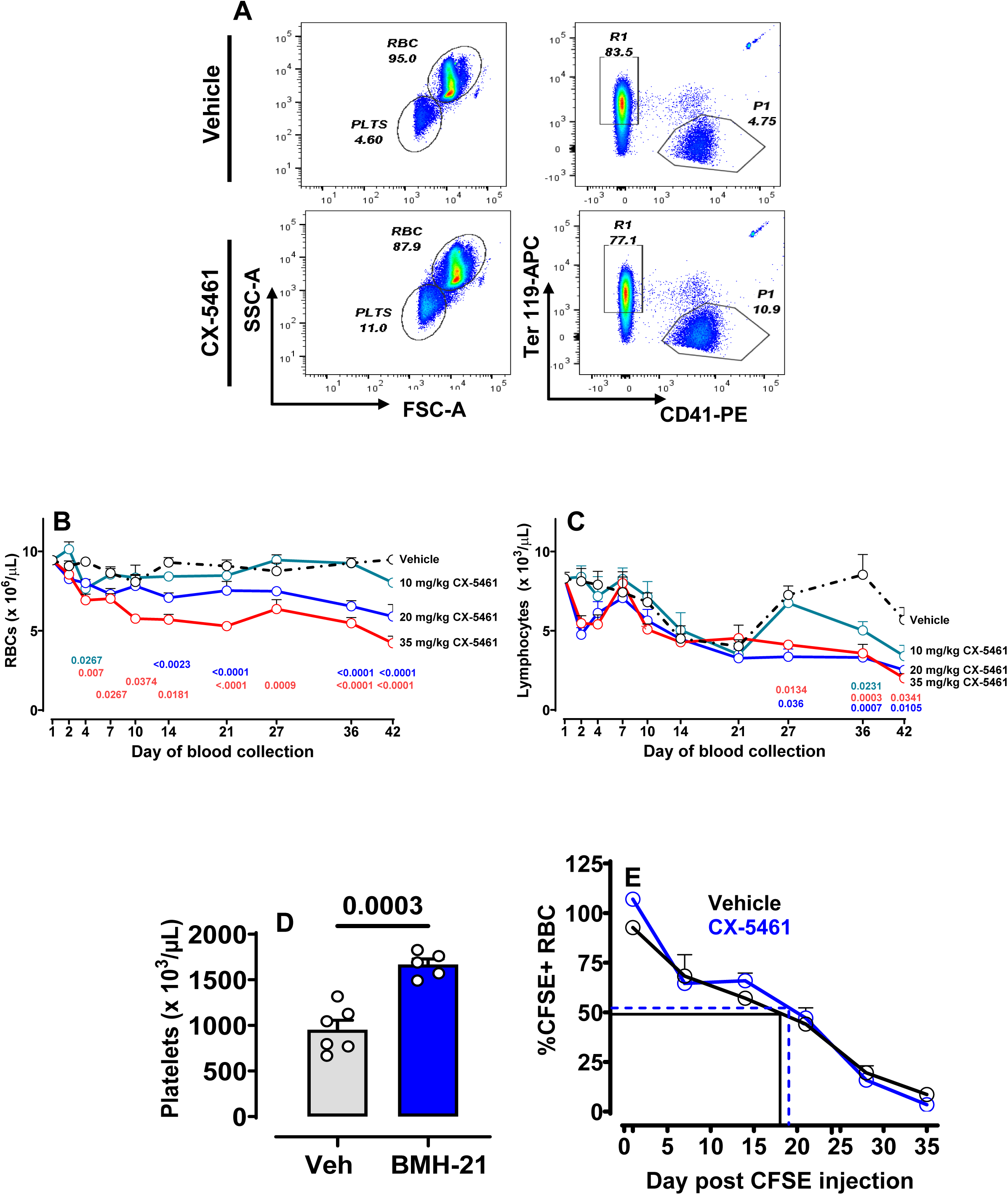
Flow cytometry approach to enumerate platelets and temporal analysis of RBC and lymphocytes in mouse treatment groups; RBC lifespan. (**A**) Gating strategy employed to quantify platelets using flow cytometry beads. C57BL/6 mice treated with 10-35 mg/kg CX-5461 for 42 days. Tail vein blood samples were collected to quantify peripheral **(B)** RBC and **(C)** WBC counts. **(D)** Peripheral platelet count in mice treated with vehicle or 72 mg/kg BMH-21. **(E)** Platelet lifespan assay: C57BL/6 mice were treated with 35 mg/kg CX-5461 or vehicle for 1 week prior to injection of CFSE *iv*. Time 0 (100%) was set at 1 h post-injection. The elimination of RBCs bearing CFSE dye over time was monitored by flow cytometry (n = 3-5 mice per timepoint) and differences in half-life were analysed. The significance among groups was analysed using Two-way ANOVA with Sidak’s multiple comparison (panels B and C) or student’s t-test (panel D and E). Statistically significant numerical p-values are presented on the graphs.

**Extended Data 3:**
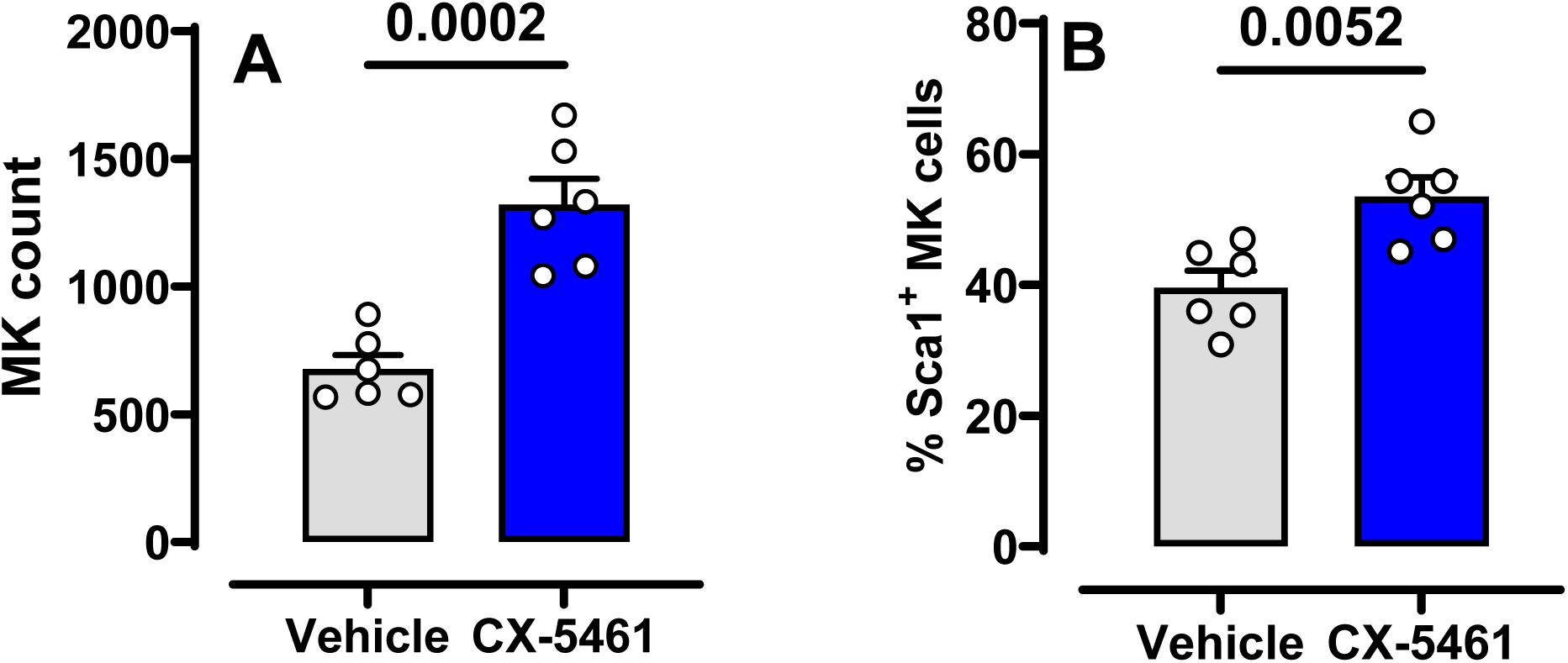
**MK levels in murine BM** BM cells were isolated from C57BL/6 mice that had been treated with 35 mg/kg CX-5461 or vehicle for 42 days and numbers of **(A)** MKs **(B)** %Sca1^+^ MKs were evaluated by flow cytometry. Data represent mean ± SEM The significance among groups was analyzed using student’s t-test. Statistically significant numerical p-values are presented on the graphs.

**Extended Data 4:**
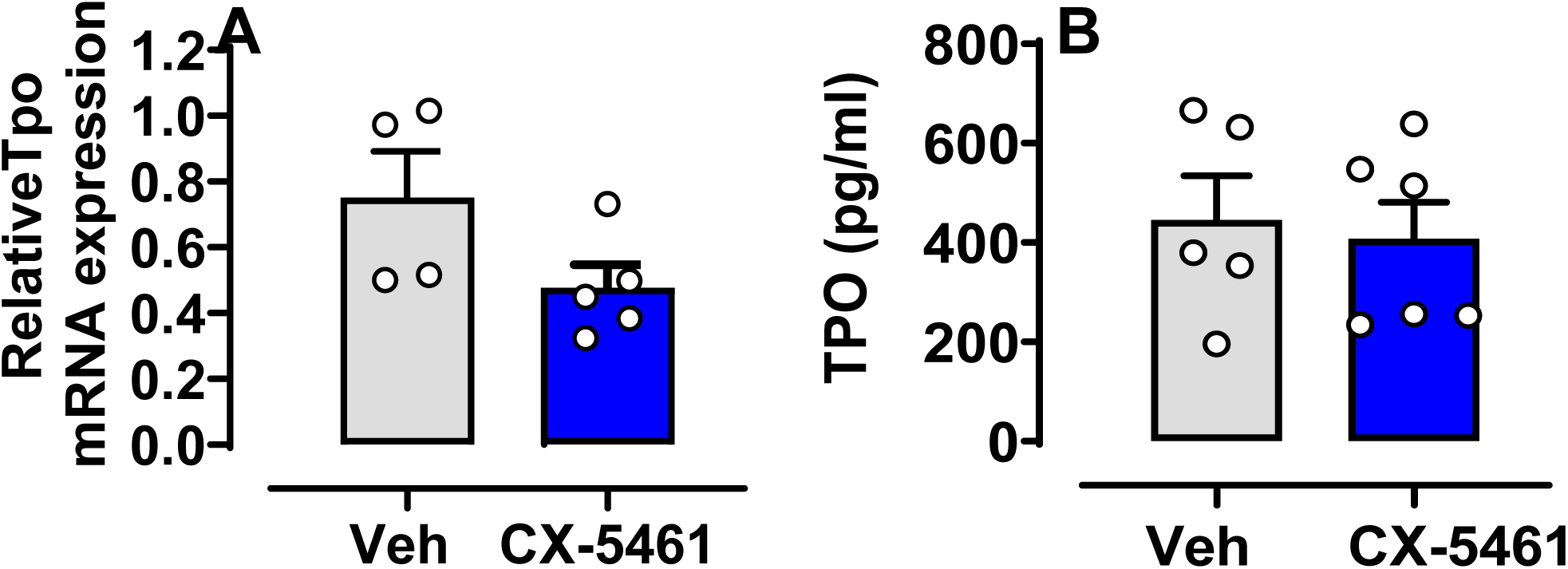
**Liver TPO mRNA levels and plasma TPO protein levels remained within normal ranges in mice treated with CX-5461.** C57BL/6 mice were treated with either 35 mg/kg CX-5461 or vehicle for 14 days and liver and plasmas were harvested. Relative levels of **(A)** liver TPO mRNA (n=4) were assessed by RT-qPCR and normalised to the phosphoglycerate kinase 1 (housekeeping) gene. **(B)** Blood from the same mice as shown in Figure 1B (treated for 42 days with either 35 mg/kg CX-5461 or vehicle) was fractionated. Plasma TPO protein was measured by ELISA. Data represent mean ± SEM. Statistical differences were analysed using student’s t-test. Statistically significant numerical p-values are presented on the graphs.

**Extended Data 5:**
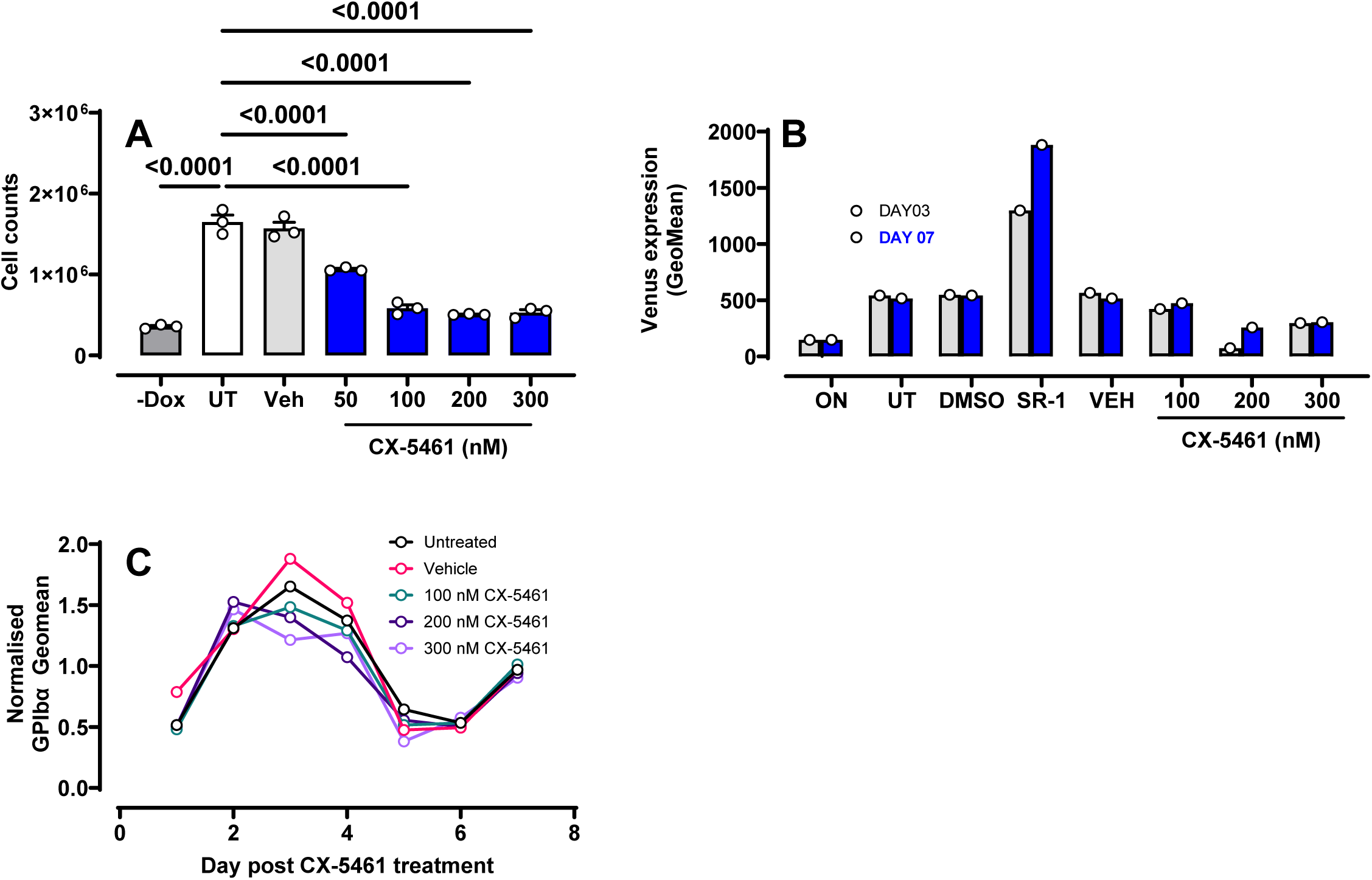
CX-5461 treatment of imMKCL inhibits cell proliferation and increases p53 expression in a dose-dependent manner. **(A)** The imMKCL line was maintained in proliferative medium containing DOX and 50-300 nM CX-5461 or an equivalent volume of vehicle. After 3 days, cell numbers were evaluated. Effect of CX-5461 on imMKCL maturation. Cells were grown under maturation conditions (without DOX) and treated with various concentrations of CX-5461 or vehicle. SR-1 treatment was used as a positive control, and cells grown under dox-on conditions as the negative control. **(B)** Venus-β1 tubulin fluorescence was measured on days 3 and 7 and **(C**) Normalised GPIbα surface expression across 7 days of treatment, which correlates to the maturation of the imMKCL cells. Data are presented as mean ± SEM and statistical significance was evaluated by one-way ANOVA with Holm-Sidak’s multiple comparison. Statistically significant numerical p-values are presented on the graphs.

**Extended Data 6:**
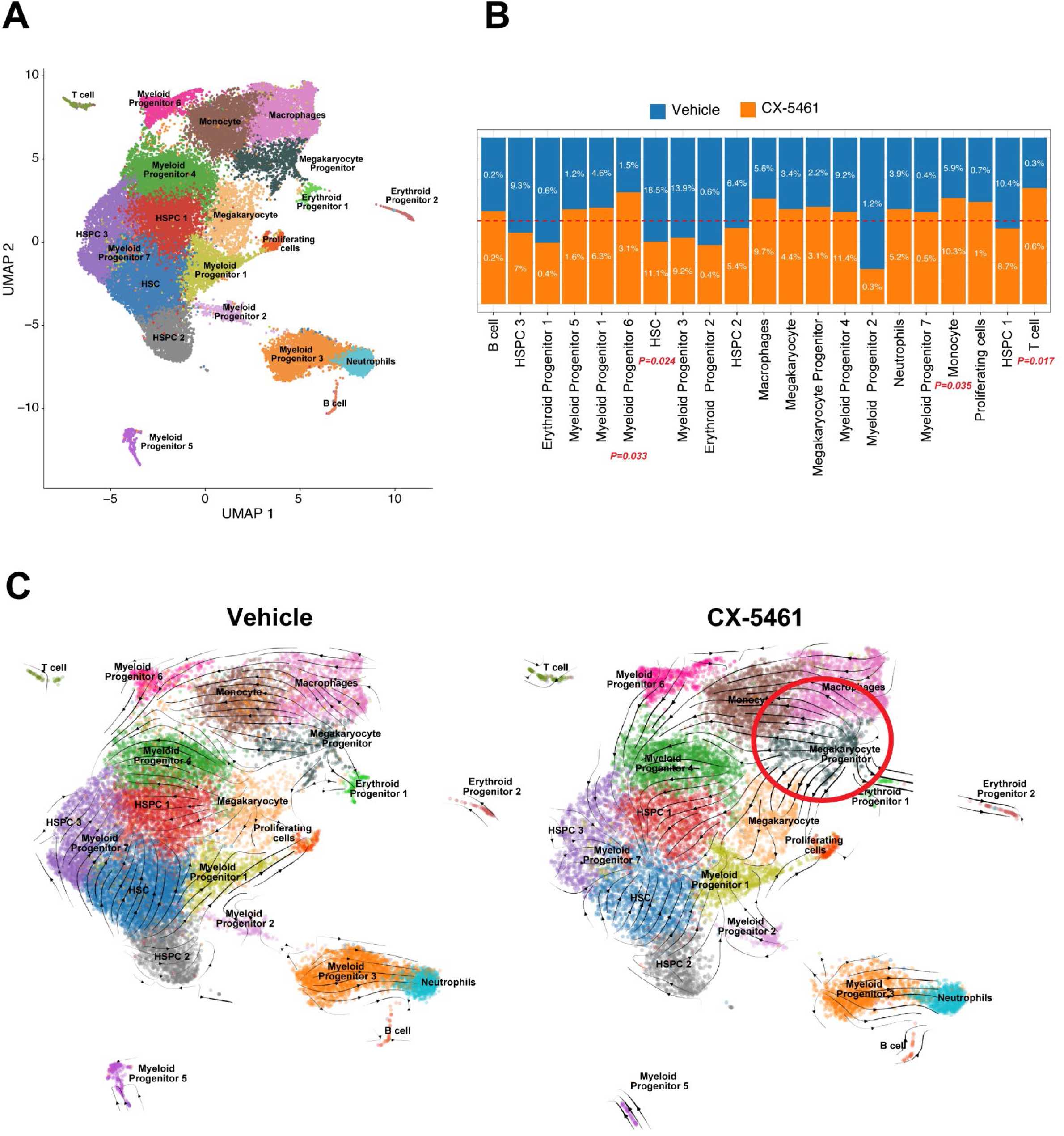
scRNA-seq analysis of mouse BM LSKs. **(A)** UMAP plots showing 21 distinct annotated clusters annotated based on known marker gene expression from pooled BM LSK cells of vehicle- and CX-5461-treated mice. **(B)** Bar plot comparing the relative frequency of each cluster between vehicle (blue) and CX-5461 (orange) treatment groups. Percentages represent the proportion of cells in each cluster out of the total number of cells per condition. Statistically significant differences in cell type proportions between treatments are indicated in red (*P*-values shown). **(C)** UMAP overlays of RNA velocity vector fields for vehicle (left) and CX-5461-treated (right) samples, illustrating inferred cell state transitions. Arrows indicate cell transition trajectories. A pronounced shift in velocity flow is observed in the CX-5461-treated group, particularly emanating from the MkP cluster (highlighted by the red circle), suggesting enhanced differentiation toward the mature megakaryocyte lineage.BM

